# Shear-Mediated Platelet Activation is Accompanied by Unique Alterations of Platelet Lipid Profile

**DOI:** 10.1101/2021.01.08.425446

**Authors:** Alice Sweedo, Lisa M. Wise, Yana Roka-Moiia, Fernando Teran Arce, S. Scott Saavedra, Jawaad Sheriff, Danny Bluestein, Marvin J. Slepian, John G. Purdy

## Abstract

Platelet activation by mechanical means such as shear stress, is a vital driver of thrombotic risk in implantable blood-contacting devices used in treatment of heart failure. Lipids are essential in platelets activation and have been studied following biochemical activation. However, little is known regarding lipid alterations occurring with mechanical – shear mediated platelet activation. Here, we determined if shear-activation of platelets induced lipidome changes that differ from those associated with biochemically-mediated platelet activation. We performed high-resolution lipidomic analysis on purified platelets from four healthy human donors. For each donor, we compared the lipidome of platelets that were non-activated or activated by shear, ADP, or thrombin treatment. We found that shear activation altered cell-associated lipids and led to the release of lipids into the extracellular environment. Shear-activated platelets released 21 phospholipids and sphingomyelins at levels statistically higher than platelets activated by biochemical stimulation. Many of the released phospholipids contained an arachidonic acid tail or were phosphatidylserine lipids, which have procoagulant properties. We conclude that shear-mediated activation of platelets alters the basal platelet lipidome. Further, these alterations differ and are unique in comparison to the lipidome of biochemically activated platelets. Our findings suggest that lipids released by shear-activated platelets may contribute to altered thrombosis in patients with implanted cardiovascular therapeutic devices.

## 1. INTRODUCTION

Thrombus formation, while valuable as a protective and reparative process, is responsible for significant morbidity and mortality in native tissue pathologies [1, 2], as well as in implanted therapeutic devices [3–6]. Central in thrombus formation is the platelet, which is activated by either biochemical or mechanical means [7, 8]. While the mechanisms of biochemical activation are well-defined, having led to the development of a wide range of anti-platelet agents in clinical use [9, 10], less is known regarding mechanisms of mechanical platelet activation.

A primary mechanical activation means of platelets is exposure to shear stress [7]. Supraphysiologic shear stress occurs when blood flow in the body is disturbed through high grade arterial or valvular stenosis. In the case of implantable cardiovascular therapeutic devices (CTD), altered blood flow is common in mechanical circulatory support devices, percutaneous heart valves, and incompletely expanded stents [11–15]. Regardless of the source of shear stress the primary interaction of shear as an activating force occurs as the platelet is subjected to accelerated, chaotic and turbulent flows [16–19]. Shear stress can impact the platelet as a whole [16, 17] or via direct impact on the lipid-rich platelet plasma membrane [18, 19].

Plasma membranes, composed mainly of lipids, are major constituents of platelets. The platelet is highly abundant in lipid-rich membrane structures. Platelets have a full internal secondary membrane system and a basal resting membrane which, upon activation, rapidly increases in surface area via contained, connected folded redundant membrane [20, 21]. In the resting state, platelet plasma membranes maintain bilayer asymmetry with sphingomyelin (SM) and phosphatidylcholine (PC) lipids on the exterior leaflet [22, 23]. Upon activation, membrane lipid reorganization occurs, associated with morphological changes and loss of membrane asymmetry leading to externalization of procoagulant lipids, including phosphatidylserine (PS) [23, 24]. The overall platelet lipidome is altered upon biochemical activation as lipids are metabolized for energy [25, 26], converted to bioactive lipids like thromboxanes involved in coagulation [26, 27], and released from internal granules [26–28]. Additionally, lipids are released in the form of prothrombotic platelet microparticles following platelet activation [25–27, 29–31]. Lipidomics is increasingly acknowledged as a method for identifying pathological pathways and biomarkers [32]. However, limited information exists regarding the effect of shear stress on the platelet lipidome.

We have previously demonstrated that biomechanical properties of platelets including overall cell stiffness [33], regional membrane stiffness [34, 35], and membrane fluidity [36, 37] are major determinants of shear-mediated platelet activation. As platelet lipids are major membrane constituents and have an overall effect on activation [20, 21], determination of the impact of shear on the composition and release of platelet lipids is vital to advance understanding of the pathophysiology of shear mediated platelet activation (SMPA). This is critically important in understanding thrombus formation in both native high shear pathologies as well as in implanted artificial devices.

We hypothesized that shear-mediated platelet activation will alter the overall platelet lipidome and the release of lipids into surrounding environment. We first tested our hypothesis by comparing the lipidome of human platelets that were non-activated or activated by shear, ADP, or thrombin treatment. Next, we examined the distribution of lipids following activation, identifying lipids retained in platelets and those released into surrounding media. Our findings suggest that lipids released by shear-activated platelets differ than those released following biochemical activation and may contribute to thrombosis in patients.

## 2. METHODS

### 2.1 Platelet purification

Platelets were collected from consenting donors (University of Arizona IRB#1810013264). Blood was collected from four adult donors. Each donor was free from aspirin or anticoagulants for at least two weeks before blood donation. Blood was collected into a vial containing Anticoagulant Citrate Dextrose Solution A via venipuncture. Platelet-rich plasma (PRP) was obtained by centrifugation at 450x g for 15 min, and further purified by column filtration (Sepharose 2B, 60-200 um diameter; Sigma-Aldrich, USA) to acquire gel-filtered platelets (GFP). The Sepharose column was pre-equilibrated with modified-Tyrode’s buffer (125mM NaCl, 10mM HEPES pH 7.4, 25mM glucose, 27mM KCl, 0.5mM Na_2_HPO_4_, 2mM MgCl_2_, 1mM sodium citrate and 0.1% bovine serum albumin (BSA)). Platelets were counted using a Z1 particle counter (Beckman Coulter, USA). Platelets were used for experiments within 2-4 h of purification.

### 2.2 Platelet activation

Each donors GFPs were divided into four sample groups: non-active (resting), shear-activated, thrombin-activated, and ADP-activated. Platelets were activated by mechanical shear, as previously described [19, 38–40]. Briefly, platelets were exposed to 70 dynes/cm^2^ shear for 10 minutes in a hemodynamic shearing device (HSD) for a shear accumulation of 42,000 dynes*s/cm^2^. These conditions model the hyper-shear forces imparted to blood components by VADs [40–42]. Activation was measured using the platelet activation state (PAS) assay [38, 42]. For biochemical activation, platelets were treated with thrombin (1 U/mL) or ADP (10 μM) in the presence of 3 mM calcium for 10 min at room temperature. Following thrombin or ADP treatment, activation was measured by flow cytometry analysis of P-selectin (CD62P) expression [43]. Non-activated control platelets were mock-treated: placed at room temperature for 10 min in buffer containing 3 mM calcium.

### 2.3 Lipid Extraction

Following activation or mock-activation, 4×10^6^ platelets were centrifuged (16,000x g for 15 min) to isolate cell-associated or released lipids (Figure 1A). The released lipids were extracted from the supernatant, while cell-associated lipids were obtained from the pellet. For lipid isolation from the supernatant fraction, 150 μl of the supernatant was transferred to a glass vial and combined with an equal volume of cold, acidic methanol for a final concentration of 50% methanol and 0.1 M HCl. Next, chloroform was added to achieve a 60:40 chloroform:methanolic solution. The samples were mixed well by gentle vortexing, before centrifugation at 1,000x g for 10 min at 4 °C to separate the liquid phases. The chloroform layer was transferred to a clean glass vial using a glass syringe. This process was repeated for a second chloroform extraction step. Lipids were dried under N_2_ gas to remove the chloroform. For lipid isolation from the platelet pellet fraction, the platelets were washed once with 1 mL ice-cold PBS prior to the addition of acidic methanol (50% methanol, 0.1 M HCl). Next, chloroform was added and lipids were extracted as described for the supernatant fraction. For each condition, lipids were extracted in parallel from 2 replicate samples. Lipids were stored at −80 °C under N_2_ gas for less than 1 month before mass spectrometry analysis. The extraction protocol was performed on platelet buffer containing no platelets to determine the background level of lipids. Chloroform (Sigma-Aldrich, USA), methanol (Fisher Chemicals, USA), and water (Fisher Chemicals, USA) used in this study were HPLC or Optima LC-MS grade.

**Figure 1.**
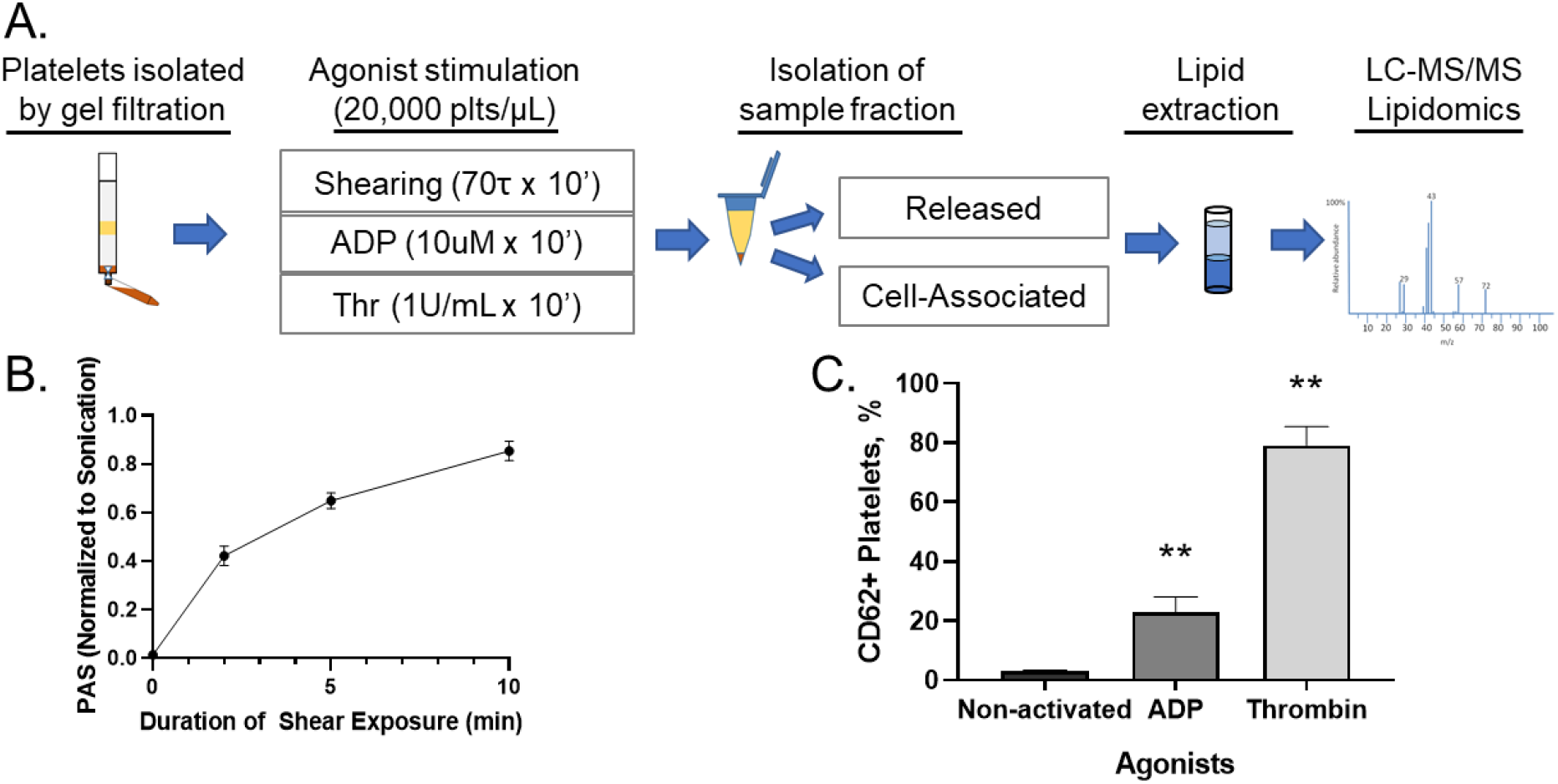
Experimental workflow and platelet activating conditions. **A.** Human platelets isolated by gel filtration were activated by either mechanical shearing or biochemical agonist (ADP or thrombin). Platelets were centrifuged and pellet (‘cell-associated) fraction was separated from the supernatant fraction (‘released). Lipids were isolated by liquid-liquid extraction and analyzed by LC-MS/MS. **B.** Platelets were activated by shear stimulus (70 dynes/cm2) in a hemodynamic shearing device (HSD). Activation level was determined by the platelet activity state (PAS) thrombogenic assay. **C.** Platelets activated by treatment with ADP (a weak agonist) and thrombin (a strong agonist). Biochemical activation was determined by flow cytometry analysis of CD62 (p-selectin) expression.

### 2.4 Mass Spectrometry

Lipids from the supernatant fraction were resuspended in 150 μL of 1:1:1 chloroform:isopropanol:methanol. Lipids from the pelleted platelets were normalized by protein concentration. Protein levels were measured using a BCA protein assay (Pierce, Thermo Scientific, USA). Lipids were resuspended in 150 μl per 35 μg of protein in 1:1:1 chloroform:isopropanol:methanol.

Lipids were identified and quantitatively measured by liquid chromatography high-resolution tandem mass spectrometry (LC-MS/MS). Lipids were separated using a Kinetex 2.6 μm C18 column (Phenomenex 00F-4462-AN) at 60 °C on a Vanquish UHPLC system (Thermo Scientific). UHPLC was performed using two solvents: solvent A (40:60 water:methanol plus 10 mM ammonium formate and 0.1% formic acid) and solvent B (10:90 methanol:isopropanol plus 10 mM ammonium formate and 0.1% formic acid) at 0.25 ml/min flow rate using conditions previously described [44]. Between each sample, the column was briefly washed and re-equilibrated. All lipids were measured using a Q-Exactive Plus mass spectrometer operating in a data-dependent Full MS/dd-MS2 TopN mode using heated electrospray ionization, as previously described.[44] Spectra were collected over a 200-1600 m/z mass range. MS1 scans were collected using a resolution setting of 70,000 or 120,000. MS2 spectra were collected using a resolution setting of 35,000. Each sample was analyzed in negative and positive modes. In negative mode, a normalized collision energy (NCE) value of 20 was used. In positive mode, the NCE value was increased to 30. The instrument was calibrated weekly.

### 2.5 Data Analysis

Lipidomic data was analyzed using MAVEN [45] and Xcalibur (Thermo Scientific) using lipid libraries from LIPIDMAPS [46]. The following lipid classes were included in the analysis: fatty acids, glycerophospholipids, sphingolipids, cholesterol esters, and glycerolipids. Guidelines from the Lipidomic Standards Initiative were followed for lipid species identification and quantification, including consideration of isotopic patterns resulting from naturally occurring ^13^C atoms and isomeric overlap. Contaminating peaks were identified by preparing samples lacking platelets. These peaks were removed from the dataset and not further considered. MS1 peaks selected in MAVEN were further analyzed to confirm their identity using MS2 fragments. The following MS2 information was used to confirm each lipid species: PC fragment of 184.074 (positive mode) and tail identification using formic adduct (negative mode), PE fragment of 196.038 or the tail plus 197.046 (negative mode) and neutral loss of 141.019 (positive mode), PG fragment of 152.996 plus the identification of the FA tails (negative mode), PI fragment of 241.012 (negative), PS neutral loss of 87.032 (negative), MG/DG/TG by neutral loss of FA tails (positive mode), CE fragment of 369.352 or neutral loss of 368.35 (positive), and sphingolipids by sphingoid backbone, e.g. 264.268 for d18:1 backbone.

### 2.6 Statistical analysis

A total of 825 unique compounds were identified in our initial lipidomics screen. Lipids included for statistical analysis were filtered using MS/MS information, and if the lipid appeared in 3 or greater donors. After filtering, a total of 265 compounds were included for statistical tests. Missing values in a sample were imputed to be the minimum value divided by 4, an imputation technique developed for datasets with missing values weighted on the lower side of the value distribution [47]. The fold change of each lipid in activated platelets relative to its level in non-activated platelets was determined for each donor. The relative fold change values were tested for statistical difference using independent t-tests. The Benjamini-Hochberg correction method [48] in the R stats package (R Core Team 2013, www.R-project.org) was used to control the false discovery rate for multiple hypothesis testing over the 265 compounds, and 0.1 q-value was deemed significant.

## 3. RESULTS

### 3.1 Platelet Activation

The platelet lipidome is complex, with 500 known and up to 5,000 putative lipid species [25, 26]. Lipids are essential for platelet function, and the lipidome changes following biochemical activation [25, 26, 30]. To determine if mechanical hyper-shear activation of platelets alters the lipidome, we isolated platelets from four healthy adult volunteers. The donors had not taken aspirin nor anticoagulants for at least two weeks before platelet isolation. Isolated platelets from each donor were separated into four groups: *i)* mock-treated (i.e., non-activated), *ii)* thrombin-treated, *iii)* ADP-treated, and *iv)* hyper-shear-treated (Figure 1A). Following activation or mock-activation, each sample was separated by centrifugation into a cellular fraction that contained platelets (i.e., “cell-associated lipids”) and a supernatant fraction that contained extracellular lipids (i.e., “released lipids”). The cell-associated and released lipids were then extracted and measured by liquid-chromatography high-resolution tandem mass spectrometry (LC-MS/MS).

Mechanical activation of platelets depends on the nature of the shear-force and duration of force experienced by platelets [19, 38–41, 43, 49]. The shear-forces experienced by cardiovascular therapeutic devices such as a ventricular assist devices (VADs) *in vivo* can be generated *in vitro* using a hemodynamic shearing device (HSD) [40, 41, 43]. We examined platelet activation as a function of time under a constant laminar shear stress environment at 70 dynes/cm^2^ using a hemodynamic shearing device (HSD) (Figure 1B). Since exposure to 10 minutes of constant 70 dynes/cm^2^ shear provided high levels of mechanically induced platelet activation (80% of maximal activation) (Figure 1B), we selected this condition for our lipidomics studies. Importantly, this HSD-treatment condition was found to be comparable to the shear stress caused by VADs [40, 41, 43]. For biochemical activation, we selected two conditions that activated platelets: 1 U/mL thrombin treatment and 10 μM ADP treatment for 10 minutes (Figure 1C). These biochemical modes of activation have been used by our group as a comparison to shear-induced activation [43].

### 3.2 Defining Lipid Diversity of Platelets from Individual Donors

First, we defined the donor-to-donor variability in the platelet lipidome following initial lipid identification by LC-MS/MS. We identified lipids in the platelet lipidome from 4 healthy donors. Our initial analysis included lipids identified in a donor’s platelets from the three modes of activation and their non-activated platelets to get a better understanding of the overall platelet lipidome for each donor. We determined the relative levels of: free fatty acids (FFA), sphingomyelin (SM), neutral lipids (NL)— cholesteryl ester (CE), monoacylglyceride (MG), diacylglyceride (DG), and triacylglyceride (TG)—and each major class of phospholipid—phosphatidylcholine (PC), phosphatidylethanolamine (PE), phosphatidylglycerol (PG), phosphatidylinositol (PI), phosphatidylserine (PS), and phosphatidic acid (PA). In total, we found peaks corresponding to 681 lipids (using a ≤5 ppm MS1 setting). While the platelet lipidome was unique to each donor, the relative ratios of these classes of lipids were similar among the four donors, e.g., PC was roughly a quarter of the total lipidome in each donor (Figure 2A and Supplemental Figure S1). Approximately half—303 lipids—were present in at least three of the four donors. From this list of 303 lipids, we generated a ‘composite platelet lipidome’ for further analysis (Figure 2B). Overall, our findings are similar to others who have found that while the platelet lipidome from each donor is unique, roughly half of identified lipids are common to most donors [25].

**Figure 2.**
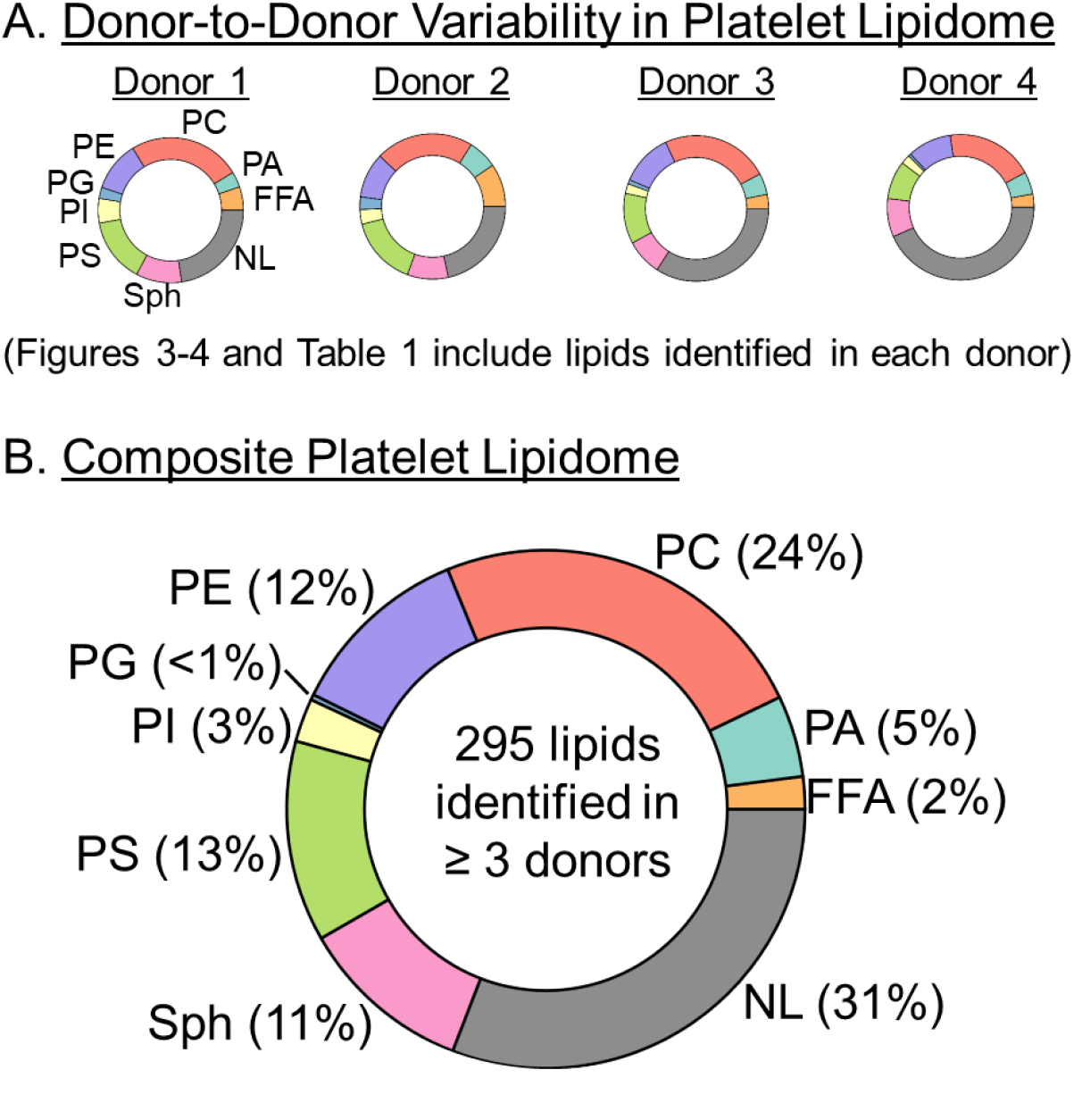
Human platelet lipidome donor-to-donor variability. **A.** The relative distribution of lipids for each of the four donors included in this study are shown by a pie chart. The data shown in Figures 3 and 4 and Table 1 represent the initial analysis of the platelet lipidome from each individual donors. **B.** A composite platelet lipidome was generated by including only lipids found in at least three of the four donors’ platelets. The relative distribution of lipids for the composite platelet lipidome is shown. The data show in Figures 5 and 6 and Table 2 represent the analyses of the lipids found in the composite lipidome shown here.

### 3.3 Shear activation of platelets alters the lipidome

Platelets can be activated by mechanical forces (e.g., shear environments) or biochemical agonists (e.g., thrombin- or ADP-containing environments). Studies of lipids in platelets have focused on biochemical activation [25, 26, 50]. We compared cell-associated lipids in HSD-treated (shear-activated) platelets to mock-treated (non-activated) platelets to determine if lipids change upon mechanical activation. Shear activation changed the platelet lipidome (Figure 3A). In this analysis, lipids in HSD-treated platelets were compared to lipids in non-activated platelets from the same donor. Several classes, including PC, PS, PE, FFA, and NL, contained lipids that were either increased or decreased following shear activation. PI and SM species tended to decrease upon shear activation.

**Figure 3.**
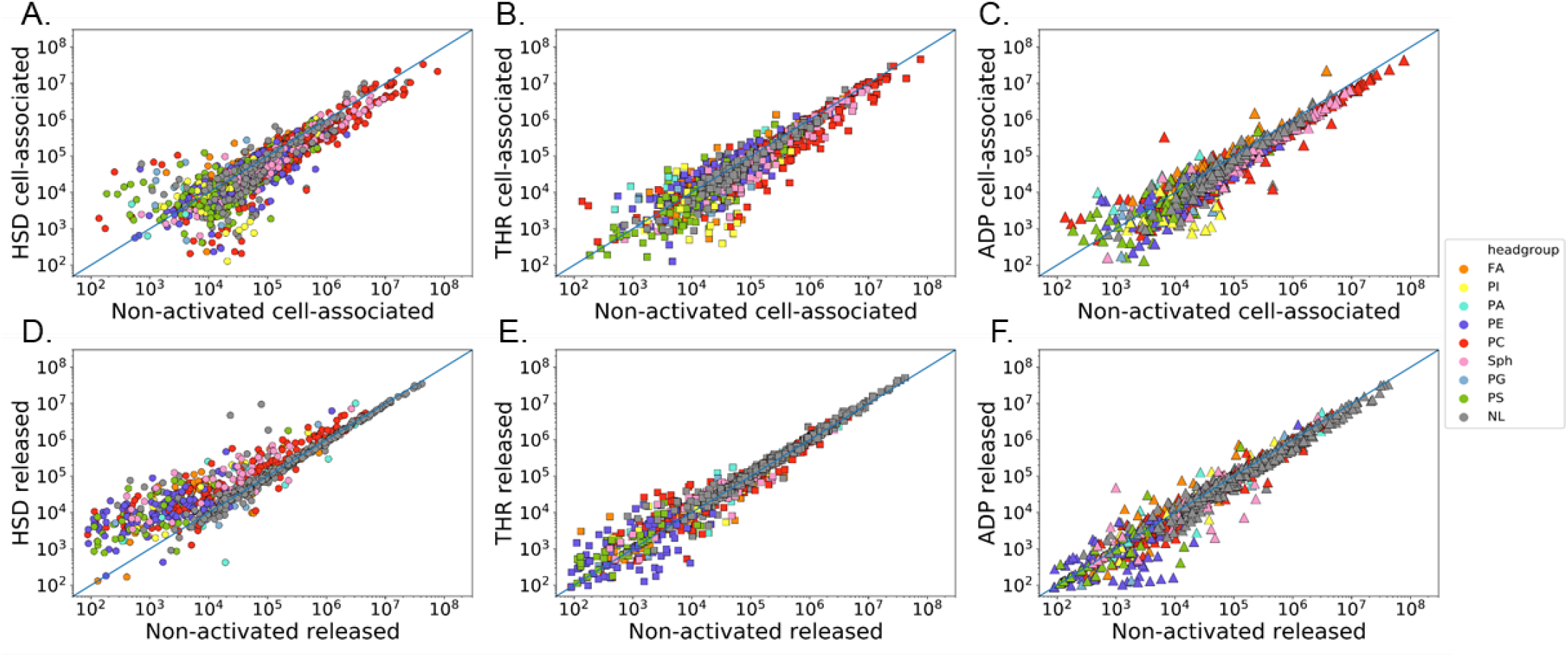
Shear activation promotes platelet lipidome changes and release of lipids. **A-C.** Relative lipid abundances of cell-associated lipids for sheared (HSD-treated), thrombin-treated (THR) and ADP-treated activated platelets (y-axis, log of ion counts), relative to non-activated (mock-treated) platelets (x-axis, log of ion counts). **D-E.** Released lipids for HSD-, THR- and ADP-treated activated platelets (y-axis, log of ion counts) compared to released lipids in non-activated control platelets (x-axis, log of ion counts). The lipids in activated platelets are set relative to the non-activated platelets from the same donor. Lipids that found at the same abundance in activated and non-activated platelets fall along the blue diagonal line.

We similarly compared lipids from biochemical activated platelets to non-activated platelets. Cell-associated lipids in several classes were changed by biochemical activation using thrombin-treatment and ADP-treatment (Figure 3B-C). Fewer lipid changes were observed in ADP-activated platelets compared to shear or thrombin activation, consistent with ADP acting as a weak-agonist for platelet activation (Figure 1C). In all three activation conditions tested, we found that the majority of lipids were unaltered in activated platelets relative to their non-activated state (Figure 3A-C).

Next, we measured lipids released by platelets into the extracellular space following shear or biochemical activation. After platelets were treated to induce activation or mock-treated, we removed the platelets and isolated the lipids released into the extracellular environment (Figure 1A). Following HSD shear activation, we observed the release of many lipid species (Figure 3D). In particular, the levels of many phospholipids and sphingomyelins (SM) were more abundant in the supernatant fraction of platelets activated by shear relative to those released by mock-treated platelets (Figure 3D). While some neutral lipids were enhanced in the supernatant fraction of shear-activated platelets, most were found to be at similar levels in shear-activated and non-activated samples. Thrombin and ADP treatments also altered the release of some phospholipids and SMs (Figure 3E-F). Like shear-treatment, most neutral lipids were unaltered by thrombin-treatment or ADP-treatment.

Notably, several lipids were found only in the supernatant fraction of HSD shear-activated platelets relative to non-activated platelets (Table 1). Although there was donor-to-donor variability in the content of lipids released, seven of these lipids contained C20:4, arachidonic acid (Table 1). These arachidonic acid-containing lipids were PE(37:5), PE(39:6), PE(39:5), PI(38:5), PS(38:5), PS(40:4), and PS(P-40:6). Our observations show that platelet activation enhances the release of lipids and suggest that shear activation induces the release of lipids differently than biochemical activation.

**Table 1.** Lipids found in the released supernatant fraction following shear activation but not observed in the released fraction of non-activated platelets.

| Compound | Tails | N (# of donors) |
| --- | --- | --- |
| LPE(22:5) | C22:5 | 3 |
| PC(O-36:0) | n.d. | 2 |
| SM(d18:0)/20:0 | n.d. | 2 |
| PE(37:5) | C20:4 + C17:1 | 1 |
| PE(39:6) | C20:4 + C19:2<sup>^</sup> / C22:5 + C17:1<sup>^</sup> | 1 |
| PE(37:3) | n.d. | 1 |
| PE(39:5) | C20:4 + C19:1 / C22:4 + C17:1<sup>^</sup> | 1 |
| PC(42:10) | n.d. | 1 |
| PC(O-40:6) | n.d. | 1 |
| PC(O-44:5) | n.d. | 1 |
| PI(38:5) | C20:4 + C18:1<sup>^</sup> | 1 |
| PS(38:5) | C18:1 + C20:4<sup>^</sup> | 1 |
| PS(40:4) | C20:4 + C18:0 | 1 |
| PS(P-40:6) | C20:4 + 18:2<sup>^</sup> / C18:1 + C20:5<sup>^</sup> / C22:4 + 16:2<sup>^</sup> | 1 |
| SM(d16:1/23:0) | n.d. | 1 |
| SM(d18:1/26:1) | n.d. | 1 |

n.d. = not determined
<sup>^</sup> These tails were unobserved in the MS/MS fragmentation analysis, and presented based on calculated FA tail composition

During activation, the morphology of platelets—including the plasma membrane—undergoes reorganization. We examined the possibility that shear activation affected the membrane integrity of platelets resulting in the movement of lipids from the cell into the extracellular environment. For each lipid species that was released, we compared their level in the released fraction relative to its level in the cell-associated fraction (Figure 4). Following shear activation, most released lipid species were found in a greater abundance in the cellular fraction than in the released fraction (Figure 4A). Thrombin- and ADP-activated platelets similarly showed that most lipids remained cell-associated (Figure 4B-C). These observations suggest that the overall membrane integrity of platelets remains intact following shear and biochemical activation [26].

**Figure 4.**
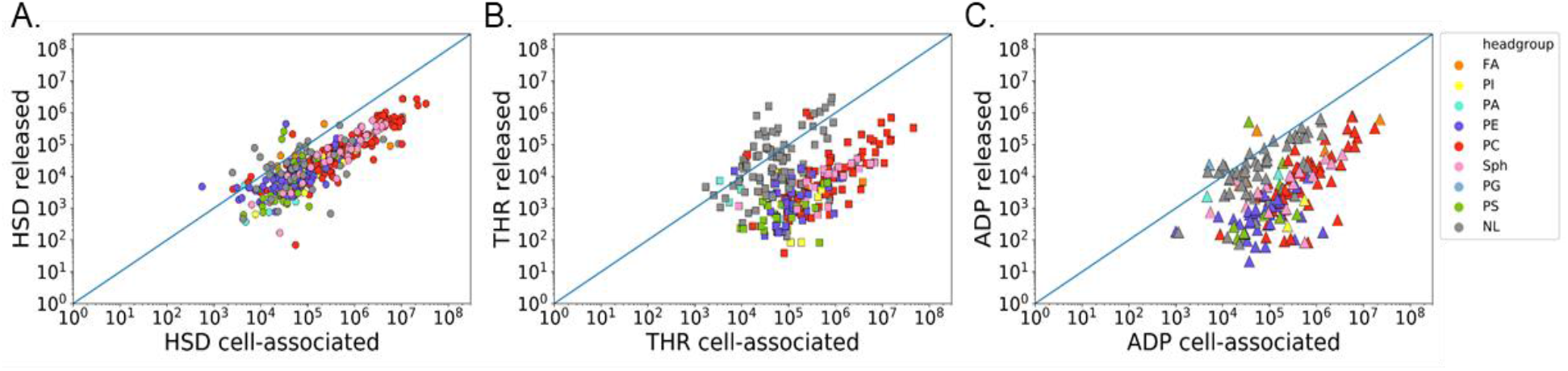
Comparison of cell-associated and released lipids following platelet activation. **A-C.** Abundance of released lipids (y-axis, log ion counts) from HSD-treated (A), THR-treated (B), and ADP-treated (C) activated platelets are compared to cell-associated lipids (x-axis, log ion counts) from the same mode of activation. Lipids below the blue diagonal line are more abundant in the cell-associated fraction, while those above the blue line are more abundant in the released fraction. Only lipids identified in both the release and cell-associated fractions are shown.

### 3.4 Shear activation promotes the release of lipids more than thrombin activation

We compared the released lipid profiles of shear- and thrombin-activated platelets to determine if the mode of activation affects the release of lipids. For this analysis, we further filtered the data to include lipids found in at least three of the four donors (i.e., the ‘composite platelet lipidome’ shown in Figure 2B). We observed that 23 of the 303 lipids were significantly different in the released fraction when comparing shear-activation and thrombin-activation (Figure 5). Most of these lipids were released at a higher level in the HSD-treated platelets compared to thrombin-treated platelets. Several phospholipids were found to be increased in the shear environment relative to thrombin, including PE, PS, and PC species. The most common tails of phospholipids were C18:1, C18:0, C16:0, and C20:4 (Table 2). We also observed C22:5, C22:4, C20:5, C20:3, and C20:1 tails in phospholipids. Several lysophosphatidylethanolamine (LPE) species and two FFAs, C18:0 and C16:0, were also significantly enhanced in the HSD shear-activated samples (Figure 5). Other FFAs, including C18:1, C18:3, and C20:5, were also enhanced in shear-activated samples compared to thrombin-treated platelets but were detectable in fewer than three donors.

**Figure 5.**
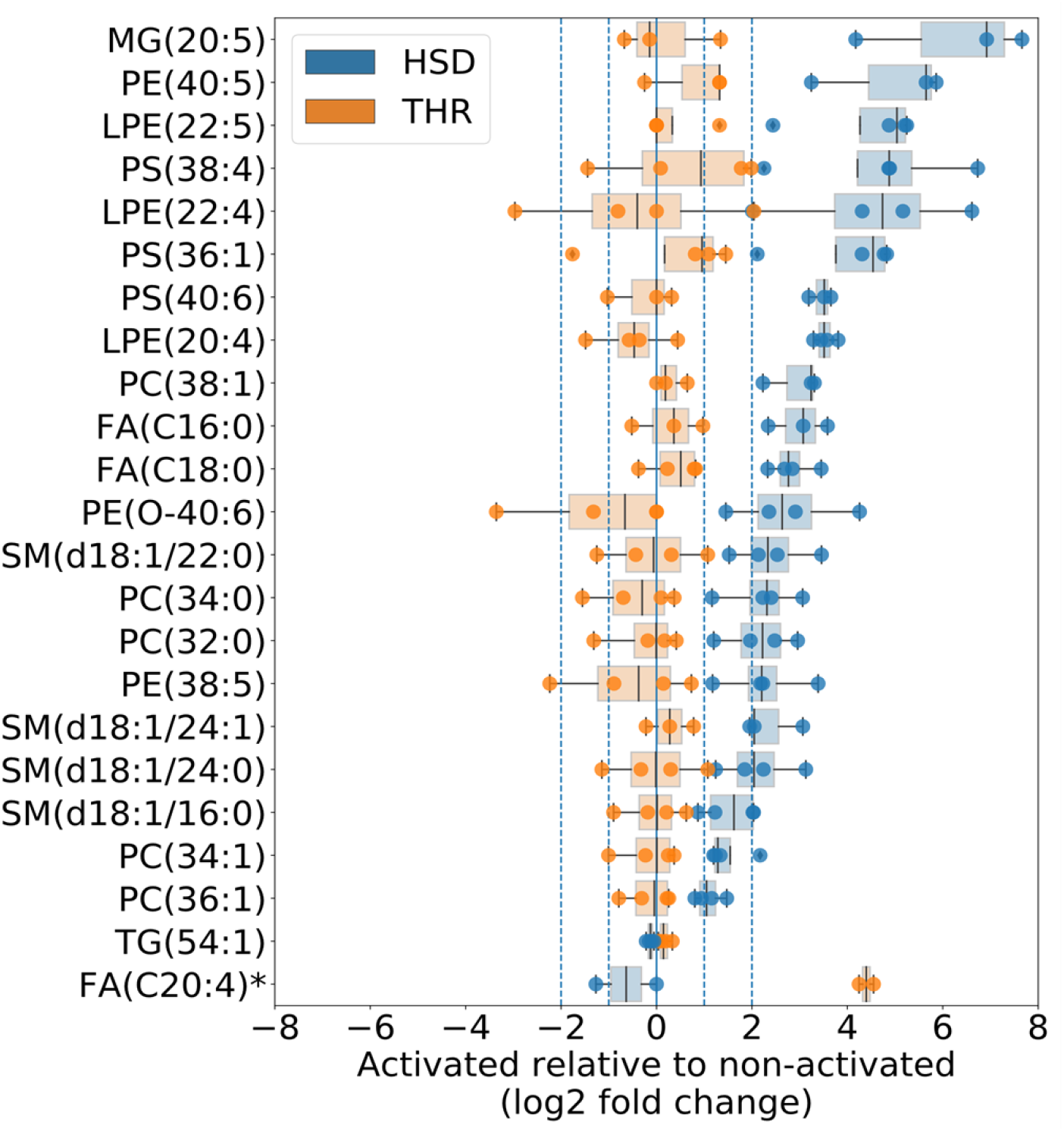
Lipids significantly different between HSD-treated shear activation and thrombin (ThR)-treated biochemical activation in the released fraction. Released lipids that are different in sheared-activated platelets compared to thrombin-activated platelets (p-value <0.1, independent t-test, corrected for multiple hypothesis testing). Lipid levels for each condition are reported as log_2_ fold change relative to non-activated controls. Each donor is represented as a dot. All lipids shown were observed in 3 or more donors, *except for Free FA(C20:4) which was only observed in 2 donors.

**Table 2.** Fatty acid composition of lipids shown in Figure 5.

|  | Fatty Acid Tails | HSD vs non<br>log2 fold<br>change<br>(released) | HSD vs non<br>log2 fold<br>change<br>(cell-associated) | HSD<br>released vs<br>HSD cell-associated |
| --- | --- | --- | --- | --- |
| **MG(20:5)** | C20:5 | 6.83 | 0.00 | inf |
| **PE(40:5)** | C20:4 + C20:1 / C22:5 + C18:0 /<br>C22:4 + C18:1 / C20:3<sup>^</sup> + C18:2 | 5.30 | -0.36 | -1.49 |
| **LPE(22:5)** | C22:5 | 4.78 | -0.71 | -1.22 |
| **PS(38:4)** | C18:0 + C20:4 / C18:1 + C20:3 /<br>C18:2 + C20:2<sup>^</sup> | 5.41 | -0.33 | -1.89 |
| **LPE(22:4)** | C22:4 | 5.30 | -0.58 | -1.55 |
| **PS(36:1)** | C18:0 + C18:1 | 4.32 | -0.43 | -1.89 |
| **PS(40:6)** | C20:4 + C20:2<sup>^</sup> / C18:0 + C22:6<sup>^</sup> | 3.47 | -0.57 | -1.29 |
| **LPE(20:4)** | C20:4 | 3.54 | -0.63 | -1.32 |
| **PC(38:1)** | C18:1 + C20:0<sup>^</sup> / C16:0 + C20:1<sup>^</sup> | 3.00 | -0.54 | -2.74 |
| **FA(C16:0)** | C16:0 | 3.09 | 0.48 | 1.08 |
| **FA(C18:0)** | C18:0 | 2.89 | 2.45 | 0.09 |
| **PE(O-40:6)** | C22:5 + C18:1<sup>^</sup> / C20:4 + C20:2<sup>^</sup> /<br>C22:4 + C18:2<sup>^</sup> | 3.11 | -0.40 | -2.43 |
| **SM(d18:1/22:0)** | C18:1 + C22:0<sup>^</sup> | 2.59 | -0.45 | -2.59 |
| **PC(34:0)** | C16:0 + C18:0 | 2.36 | -0.45 | -2.87 |
| **PC(32:0)** | C16:0 + C16:0 | 2.29 | -0.49 | -2.58 |
| **PE(38:5)** | C20:4 + C18:1 / C22:5 + C16:0 /<br>C20:5 + C18:0 | 2.46 | -0.31 | 2.03 |
| **SM(d18:1/24:1)** | C18:1 + C24:1<sup>^</sup> | 2.45 | -0.54 | -2.28 |
| **SM(d18:1/24:0)** | C18:1 + C24:0<sup>^</sup> | 2.28 | -0.62 | -2.41 |
| **SM(d18:1/16:0)** | C18:1 + C16:0<sup>^</sup> | 1.62 | -0.68 | -1.16 |
| **PC(34:1)** | C18:1 + C16:0 | 1.55 | -0.50 | -2.21 |
| **PC(36:1)** | C18:1 + C18:0 / C16:0 + C20:1 | 1.12 | -0.56 | -2.20 |
| **TG(54:1)** | C16:0 + C20:0 + C18:1<sup>^</sup> | -0.13 | -0.64 | 2.50 |

<sup>^</sup> These tails were unobserved in the MS/MS fragmentation analysis, and presented based on calculated FA tail composition

Next, we examined if the lipids in Figure 5 were different in the cell-associated lipid fraction. The same type of analysis found that there was no difference between these lipids in the cell-associated fractions (Supplemental Figure S2). For both thrombin- and shear-activated platelets, MG(20:5) was only detectable in the supernatant fraction and was not observed in the cell-associated fraction. These observations show that while the two modes of platelet activation have similar effects on cell-associated lipid changes, the release of lipids from the activated platelets differ.

In contrast to the other lipids listed in Figure 5, we observed a higher level of free C20:4 (arachidonic acid) released in thrombin-activated platelets than shear-activated platelets (Figure 5). Relative to non-activated platelets, arachidonic acid was 16-fold higher in the supernatant fraction of thrombin-treated platelets. In the shear-activated platelets of two donors, the level of released free arachidonic acid was similar to that observed in non-activated platelets. We were unable to observe free C20:4 in the other two donors, suggesting that the release of arachidonic acid following activation varies from person-to-person.

Overall, our data demonstrate that platelet activation by HSD-treatment and thrombin-treatment induce the release of lipids. However, shear activation promotes the release of lipids more so than biochemical activation.

### 3.5 Shear activation alters the platelet lipidome in ways still yet to be defined

We found a lipid that was >16-fold higher in the supernatant of shear-activated platelets relative to non-activated or thrombin-activated platelets (Figure 6A-B). This lipid has a mass-to-charge ratio (m/z) of 782.57 at z=-1, which corresponds to PC(36:3) and PE(39:3). However, we could not confirm either one of these identifications based on MS/MS fragmentation. Instead, we observed fragments corresponding to C20:4 and C22:4 fatty acyl tails (Figure 6C and Supplemental S3). Although the identification of this lipid remains unknown, it nonetheless highlights that mechanical activation of platelets creates unique lipid changes when compared to activation by biochemical agonists in ways still unknown.

**Figure 6.**
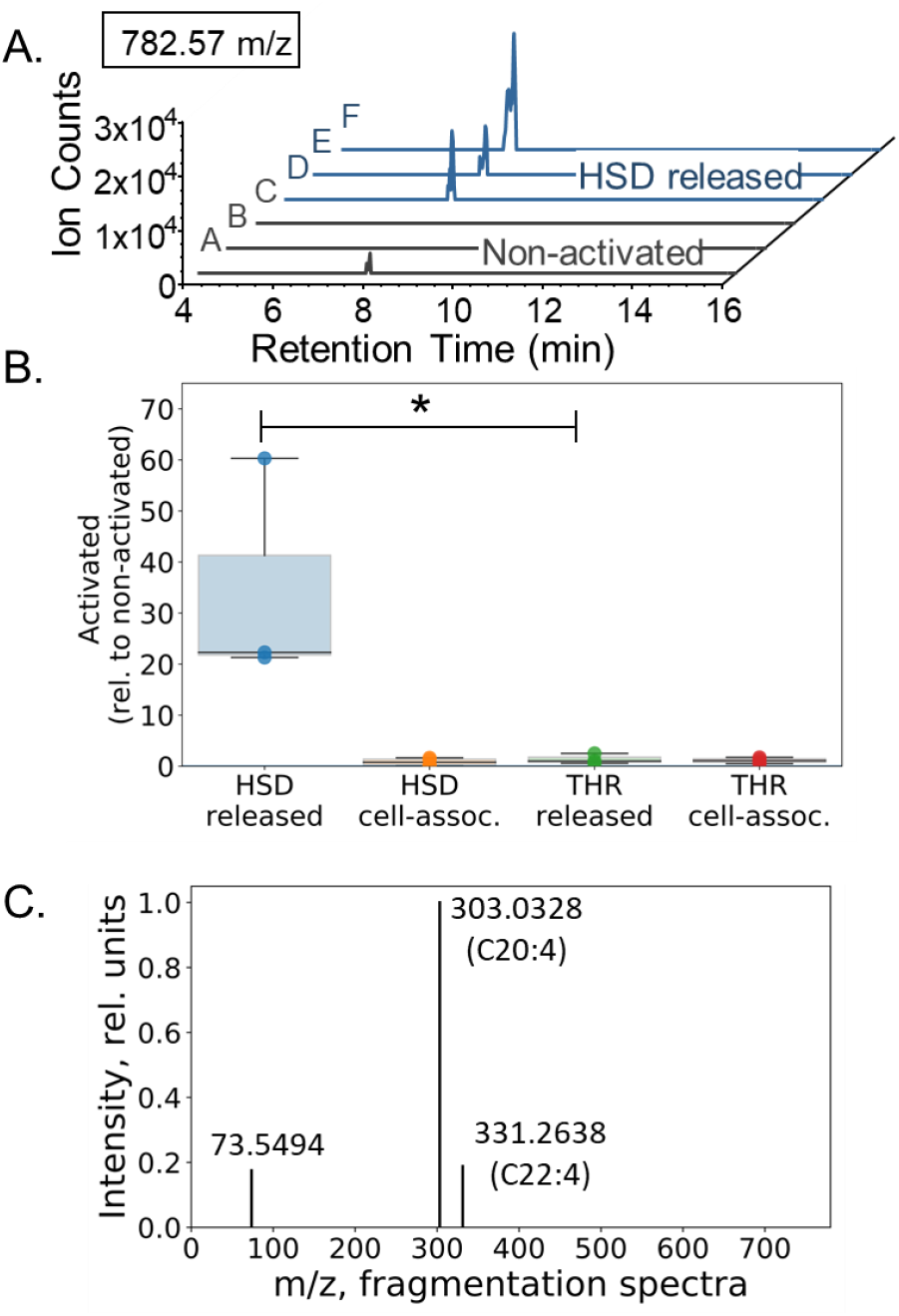
An unidentified lipid was observed in the released fraction of HSD-treated shear activated platelets. **A.** A lipid of m/z 782.57 (z=-1) was detected in the released fraction of shear-activated platelets, but not in non-activated control platelets. **B.** Lipid 782.57 is upregulated only in the HSD-treated released fraction, and not in THR-treated released or cell-associated fractions. **C.** Common MS/MS fragments observed for lipid 782.57. Masses for C20:4 and C22:4 FA tails are labeled.

## 4. DISCUSSION

Currently over 6 million people are diagnosed with heart failure, and an estimated 50% increase in prevalence is predicted in the next 10 years [48]. As a result of the rise in heart failure, the use of ventricular assist devices that promote blood flow will increase. While advancements in VAD design have reduced shear stress, VADs are still a significant source of hyper-shear forces [15], leading to shear-induced thrombosis [51]. As such, it is essential to understand the mechanical activation of platelets and how it may differ from other modes of activation. This study investigated the alteration of the platelet lipidome following shear-activation. We found that mechanical activation led to several changes in the platelet lipidome, including the release of lipids into the extracellular environment. While studies have previously examined the effect of select biochemical agonists on platelet lipidomics [25–27, 30, 31], this is the first study to examine the lipid-modulating impact of a mechanical agonist. Importantly, we found that mechanical activation creates a unique lipidome that is different from that of platelets activated by thrombin or ADP.

Activated platelets participate in the process of coagulation via aggregation and by supplying lipids, which act to regulate coagulant activities. The phospholipid headgroup is an important attribute in determining a lipid’s procoagulant activity [52]. Following platelet activation, PS lipids—which are typically sequestered on the interior of the platelet—are externalized and support the binding and assembly of coagulation factors that promote thrombin generation [24, 30, 53–55]. We previously established that PS externalization in platelets is a marker of shear activation [43]. In this study, we found that PS lipids were significantly elevated in the released fraction of shear-activated HSD-treated platelets (Table 1 and Figure 5). Specifically, we observed that PS(36:1), PS(38:4), and PS(40:6) were released by shear-activated platelets at a higher level than thrombin treatment. Since PS lipids enhance coagulation following thrombin treatment [30], our observations of high levels of PS release by HSD-treated platelets suggest that lipids released upon shear activation may enhance coagulation similar to or greater than thrombin.

PI lipids are considered to be procoagulant, though weaker than PS [56]. In one donor, PI(38:5) containing an arachidonic acid tail was released following shear treatment (Table 1). Furthermore, PI, PE, PG, and PA act synergistically with PS to enhance coagulation [30, 52]. While several PI, PG, and PA lipids were released following shear activation, these lipids were similar in the supernatant fraction of shear-activated platelets and thrombin-activated platelets, in general (Figure. 3D). However, some PE, LPE, and PC lipids were released in HSD-treated platelets at levels higher than in platelets treated with thrombin (Table 2, Figure 5), suggesting that they may be related coagulation in a shear environment.

Arachidonic acid (C20:4) liberated from membrane phospholipids is a significant driver in platelet activation by biochemical agonists such as thrombin, collagen, and ADP. Arachidonic acid can be metabolized by cyclooxygenases (COX-1 and COX-2) in activated platelets. Aspirin inhibition of COX-1 and COX-2 reduces biochemical activation of platelets, however up to 20% of patients taking aspirin due to a previous thrombotic event will have a recurrent thrombotic event [57]. Aspirin-resistant platelet activation may occur through COX-independent pathways [57–59]. The role of arachidonic acid in shear-mediated activation is not as well studied, but it is noteworthy that aspirin is ineffective in preventing shear-mediated platelet activation [38, 39]. We observed PI(38:5), PS(38:5), PS(40:4), PS(P-40:6), and several PE species containing an arachidonic acid tail were differentially affected by HSD-treatment and thrombin-treatment (Figure 5 and Table 2). For example, we observed that PS(38:4) with an arachidonic acid tail, a lipid with known procoagulant activity [30], was significantly elevated in shear-activated platelets. In some of these cases, the arachidonic acid-containing lipid was observed in fewer than three donors, suggesting that future studies with larger populations will likely yield additional findings related to possible procoagulant lipids released by mechanically-activated platelets. Additionally, the 782.57m/z unidentified lipid that was released following HSD-treatment but not thrombin-treatment also contained an arachidonic acid tail (Figure 6). Our observations that lipids with arachidonic acid differ in shear-activated and biochemically-activated platelets suggest that lipid metabolism may contribute to the aspirin resistance observed in shear-activated platelets.

In contrast to the procoagulant properties of some phospholipids, SM may have anticoagulant properties [60]. While we found several SM lipids were released at a higher level in shear-activated platelets relative to thrombin-activated cells (Figure 5), it is currently unknown if these specific SM lipids alter coagulation. Additionally, some fatty acids, such as C20:5 (eicosapentaenoic acid) and C22:5 (docosapentaenoic acid), reduce the procoagulant effects of arachidonic acid [61–63]. Several lipids with C20:5 and C22:5 tails were enhanced by shear-activation (Table 2). However, we did not observe the free fatty acid form, suggesting that their ability to affect coagulation may be limited.

Shear-treatment of platelets generates microparticles, which are also observed in platelets activated by thrombin and calcium ionophore A23187 [28, 30, 55, 64]. The role of these microparticles in thrombosis remains to be defined, though evidence suggests that shear-derived microparticles may be procoagulant, similar to those shed from biochemically activated platelets [30, 55]. Since we observed significant differences in the lipids released in HSD-treated platelets relative to thrombin-treatment, our findings further suggest that shear and biochemical activation induce different types of microparticles or mechanisms to generate microparticles. The lipidomic observations presented in this study indicate that further consideration should be placed on determining how different modes of platelet activation may affect the procoagulant properties of released microparticles.

Lipid composition contributes to the chemical and mechanical properties of platelets. Defining the platelet lipidome is important to understand how lipids function before and after activation. Although platelets have donor-to-donor variability, a lipid profile common to platelets from donors can be defined (Figure 2 and Supplemental Figure S1). Our findings were consistent with a previous study that found platelets from three donors had approximately 65% of lipids in common [25]. In this study, we found an unidentified lipid increased by shear activation, 782.57 m/z (Figure 6 and Supplemental Figure S3). This lipid species was unaltered by thrombin or ADP treatment. Slatter, et al., observed a lipid with a similar mass, 782.5691 m/z, in an untargeted lipidomics analysis [25]. The previous work found that this lipid was unchanged by thrombin treatment, similar to our observation. These observations highlight the need to continue to examine lipids using state-of-the-art approaches and various models of activation.

In conclusion, by comparing the platelet lipidome before and after activation by mechanical and biochemical stimuli, we show that shear exposure leads to a lipid profile different than biochemically-activated and non-activated platelets. While changes were found in cell-associated lipids, lipidomic differences were most evident in the lipids released into the extracellular space. Our findings highlight the relevance of lipid contribution to platelet function and the continued need to understand platelet activation under hyper-physiological shear conditions created by cardiovascular therapeutic devices such as VADs in patients with heart failure. Overall, our findings demonstrate that hyper-shear activation of platelets induces unique lipid changes and may contribute to altered thrombogenic potential of platelets in patients with implanted VADs.

## Abbreviations

BSA: bovine serum albumin;
CE: cholesteryl ester lipids;
CTD: cardiovascular therapeutic devices;
DG: diacylglycerol lipids;
FFA: free fatty acid;
GFP: gel-filtered platelets;
HSD: hemodynamic shearing device;
LC-MS/MS: liquid chromatography high-resolution tandem mass spectrometry;
LPE: lysophosphatidylethanolamine lipids;
MG: monoacylglycerol lipids;
m/z: mass-to-charge ratio;
NCE: normalized collision energy;
NL: neutral lipids;
PAS: platelet activation state assay;
PA: phosphatidic acid lipids;
PC: phosphatidylcholine lipids;
PE: phosphatidylethanolamine lipids;
PG: phosphatidylglycerol lipids;
PI: phosphatidylinositol lipids;
PRP: platelet-rich plasma;
PS: phosphatidylserine lipids;
SM: sphingomyelin lipids;
SMPA: shear mediated platelet activation;
TG: triacylglycerol lipids;
VADs: ventricular assist devices

## CRediT AUTHORSHIP CONTRIBUTION STATEMENT

**Alice Sweedo:** Conceptualization, Data curation, Formal Analysis, Investigation, Methodology, Visualization, Validation, Writing – original draft, Writing – review & editing. **Lisa M. Wise:** Conceptualization, Data curation, Formal Analysis, Investigation, Methodology, Visualization, Validation, Writing – original draft, Writing – review & editing. **Yana Roka-Moii:** Data curation, Formal Analysis, Investigation, Writing – review & editing. **Fernando Teran Arce:** Conceptualization, Writing – review & editing. **S. Scott Saavedra:** Conceptualization, Funding Acquisition, Writing – review & editing. **Jawaad Sheriff:** Conceptualization, Funding Acquisition, Writing – review & editing. **Danny Bluestein:** Conceptualization, Funding Acquisition, Writing – review & editing. **Marvin J. Slepian:** Conceptualization, Funding Acquisition, Project administration, Supervision, Writing – original draft, Writing – review & editing. **John G. Purdy:** Conceptualization, Data curation, Formal analysis, Funding Acquisition, Project administration, Supervision, Writing – original draft, Writing – review & editing.

## FUNDING

This work was supported by the Arizona Biomedical Research Commission through the Arizona Department of Health Services (ADHS18-198868 to J.G.P.), the National Institute of Health (NIH) (T32HL007955 to A.S. and 5U01HL131052 to D.B. and M.J.S.), BIO5 Institute at the University of Arizona (Pilot Interdisciplinary Project to J.G.P, S.S.S. and M.J.S.). The content is solely the responsibility of the authors and does not necessarily represent the views of ADHS and NIH.

## DECLARATION OF INTEREST

Declarations of interest: none.

## ACKNOWLEDGMENTS

We thank Debbie Mustacich for assistance in the initial stages of this project. We also thank the University of Arizona BIO5 Institute Statistics Consulting Lab for helpful discussion.

---

**Figure Supplemental S1.**
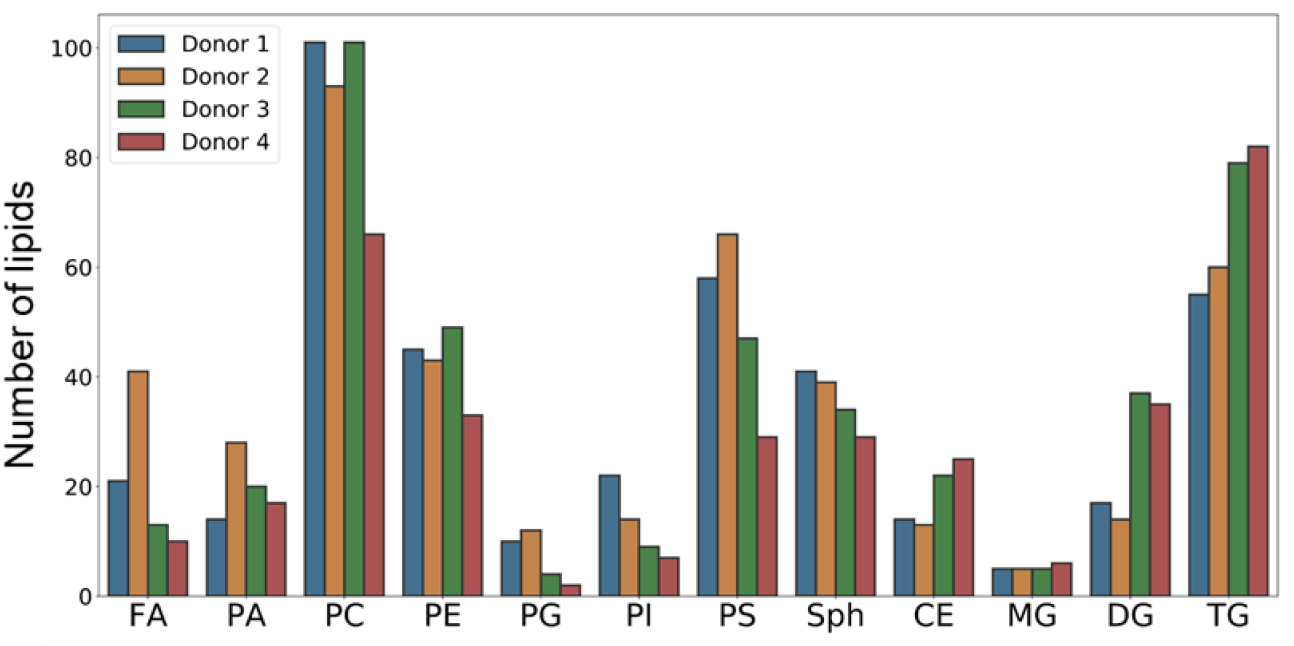
Platelet lipidomes from individual donors. The number of lipids identified by MS1 analysis are shown for each class of lipids.

**Figure Supplemental S2.**
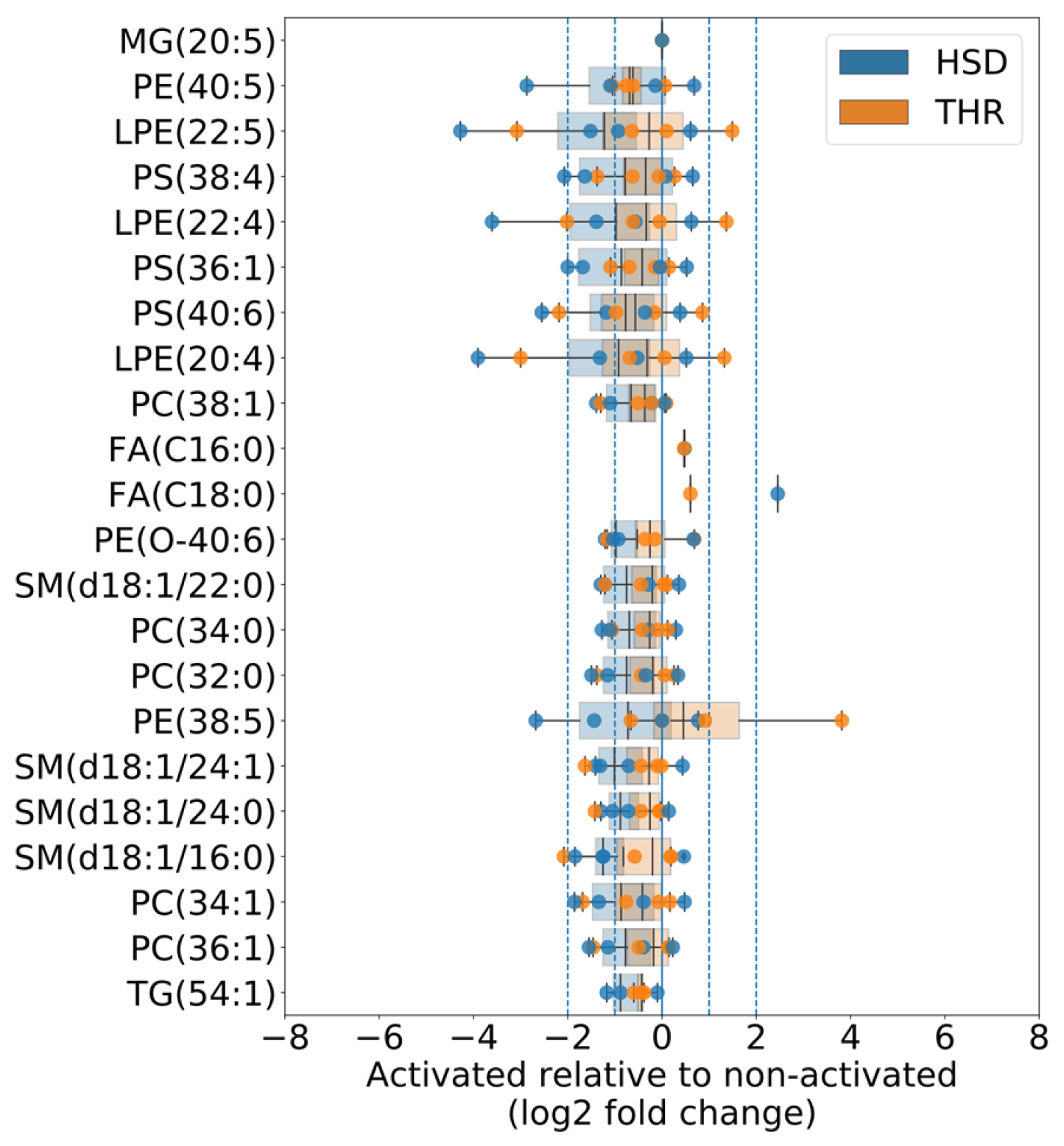
Analysis of cell-associated fraction of lipids shown in Figure 5. MG(20:5) and Free FA(C20:4) were not observed in the cell-associated fraction.

**Figure Supplemental S3.**
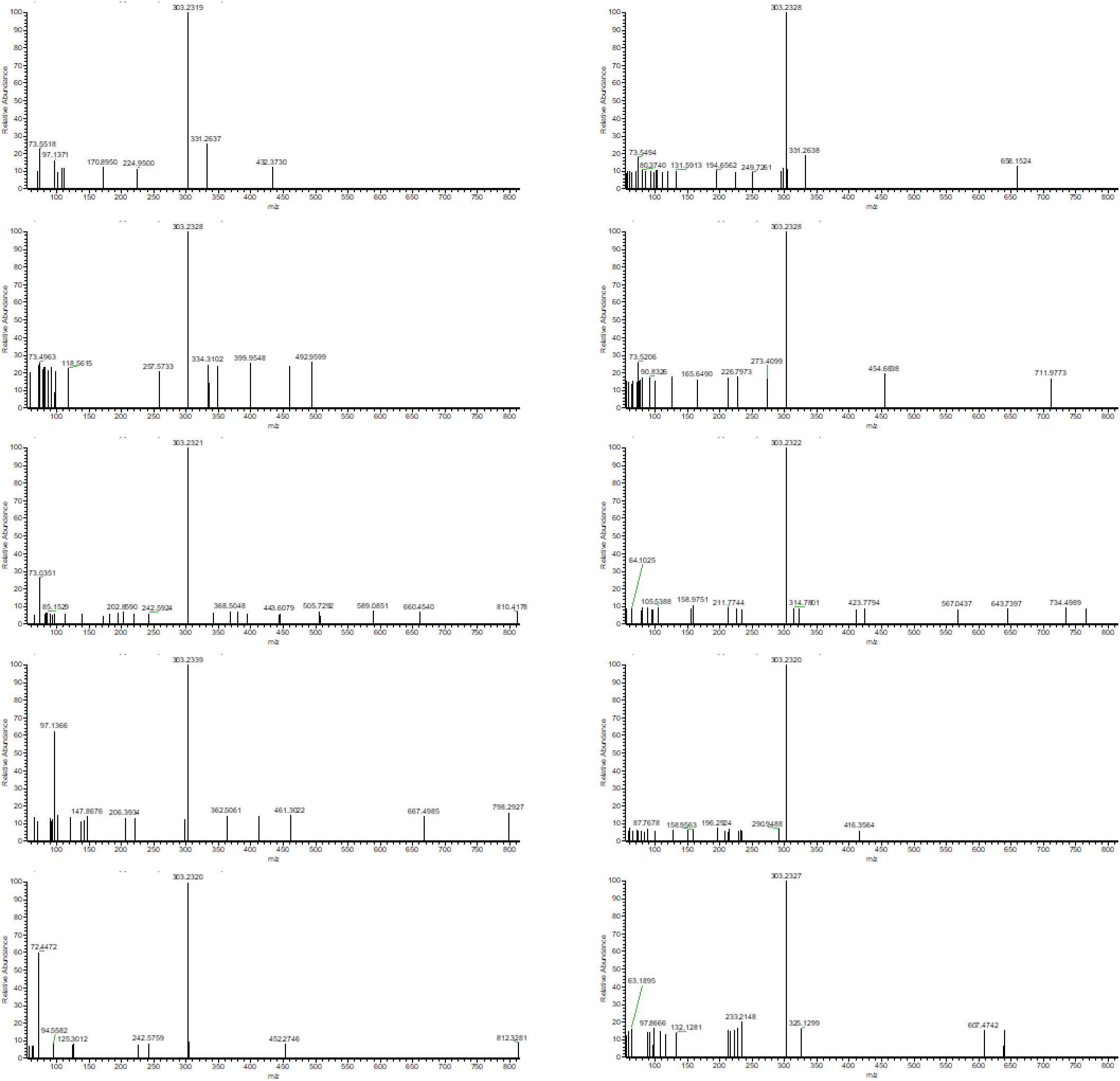
MS/MS fragment spectra of unknown lipid 782.57 shown in Figure 6.

## Notes

### Competing Interest Statement

The authors have declared no competing interest.

